# The adapted Activity-By-Contact model for enhancer-gene assignment and its application to single-cell data

**DOI:** 10.1101/2022.01.28.478202

**Authors:** Dennis Hecker, Fatemeh Behjati Ardakani, Alexander Karollus, Julien Gagneur, Marcel H. Schulz

## Abstract

Identifying regulatory regions in the genome is of great interest for understanding the epigenomic landscape in cells. One fundamental challenge in this context is to find the target genes whose expression is affected by the regulatory regions. A recent successful method is the Activity-By-Contact (ABC) model (Fulco et al., 2019) which scores enhancer-gene interactions based on enhancer activity and the contact frequency of an enhancer to its target gene. However, it describes regulatory interactions entirely from a gene’s perspective, and does not account for all the candidate target genes of an enhancer. In addition, the ABC-model requires two types of assays to measure enhancer activity, which limits the applicability. Moreover, there is no implementation available that could allow for an integration with transcription factor (TF) binding information nor an efficient analysis of single-cell data. We demonstrate that the ABC-score can yield a higher accuracy by adapting the enhancer activity according to the number of contacts the enhancer has to its candidate target genes and also by considering all annotated transcription start sites of a gene. Further, we show that the model is comparably accurate with only one assay to measure enhancer activity. We combined our generalised ABC-model (gABC) with TF binding information and illustrate an analysis of a single-cell ATAC-seq data set of the human heart, where we were able to characterise cell type-specific regulatory interactions and predict gene expression based on transcription factor affinities. All executed processing steps are incorporated into our new computational pipeline STARE. The software is available at https://github.com/schulzlab/STARE.

## 1 Introduction

Unravelling the mechanisms behind gene expression regulation is a central task in epigenomics. Enhancers are key players in this process. They are accessible regions in the genome, which can be bound by transcription factors (TFs) in a sequence-specific fashion. Those TFs have a variety of functions: recruit other cofactors, remodel chromatin conformation, cause changes in epigenetic modifications or directly interact with the transcription machinery, affecting gene expression (Pabo and Sauer, 1992; Gonzalez, 2016; Lambert et al., 2018). Many methods exist for annotating enhancers, *e*.*g*., using open-chromatin assays like DNase-, ATAC- or NOMe-seq (Song and Crawford, 2010; Buenrostro et al., 2015; Kelly et al., 2012). Histone modifications associated with enhancer activity like H3K27ac or H3K4me1 also aid enhancer annotation (Heintzman et al., 2009; Creyghton et al., 2010). Besides a plethora of methods to define enhancers, another ongoing challenge is to identify target genes of enhancers, which is essential for understanding their function. These enhancer-gene interactions can span large distances and are insufficiently explained by linear proximity (Yao et al., 2015; Schoenfelder and Fraser, 2019). Many approaches exist to predict target genes of enhancers, *e*.*g*., using correlation of enhancer activity and gene expression (The FANTOM Consortium et al., 2014; Gao and Qian, 2019; Schmidt et al., 2021), or correlation of accessibility across samples (Pliner et al., 2018). Another possibility is to include chromatin contact data, *e*.*g*. Hi-C (Lieberman-Aiden et al., 2009), to call chromatin loops for annotating enhancer-gene links (Rao et al., 2014; Schmidt et al., 2020; Yi et al., 2021). Although loops correlate with gene expression, their anchors only cover a fraction of active promoters and enhancers and their removal impacts expression of only few genes (Nora et al., 2017; Rao et al., 2017; Schoenfelder and Fraser, 2019). Fulco et al. (2019) combined measurements of enhancer activity with chromatin contact data and proposed the Activity-By-Contact (ABC) model. The assumption is that active enhancers that frequently contact a gene’s promoter are more likely to affect a gene’s regulation. The ABC-model requires DNase-seq, H3K27ac ChIP-seq data and a Hi-C matrix. The Hi-C matrix can be substituted with a matrix averaged over multiple cell types, or with a quantitative function describing the distance-contact relationship (Fulco et al., 2019). The original ABC-model formulation is entirely gene-centric, which means it does not take the candidate target genes of an enhancer into account.

We propose a generalised ABC-score with two adaptations: firstly it describes enhancer activity in a gene-specific manner, and secondly it uses the information of all annotated transcription start sites (TSSs) of a gene. Further, we could show that, instead of using both DNase- and H3K27ac ChIP-seq data for the ABC-model, one assay for measuring enhancer activity yields a similar accuracy. We validated the adaptations on three data sets of experimentally tested enhancer-gene interactions in K562 cells and on expression quantitative trait loci (eQTL) data from different tissues. We developed STARE, a fast implementation of the ABC-score with an approach to quantify TF binding affinity for genes. STARE can compute enhancer-gene interactions from single-cell chromatin accessibility data, illustrated on data of the human heart. With only one data modality at single-cell resolution, we identified and characterised cell type-specific regulatory interactions.

## 2 Methods

### 2.1 Activity-By-Contact score

We use the terms enhancer and regulatory region interchangeably. The principle of the ABC-model is that an enhancer, which is highly active and has a high contact frequency with a gene, is likely to regulate it (Fulco et al., 2019). The ABC-score represents the relative contribution of an enhancer *r* to the regulation of gene *g*, measured by the enhancer’s activity *A*_*r*_ and its contact frequency *C*_*r,g*_ with the promoter of *g*. For each candidate enhancer the activity is multiplied with the contact frequency and this product is taken relative to the sum over all candidate enhancers *R*_*g*_ in a predefined window around the TSS of a gene:

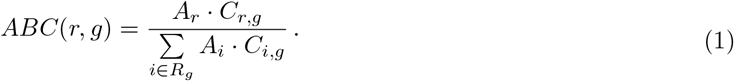

By definition the scores per gene sum up to 1. In practice, a cut-off is used to select valid interactions. The ABC-model allows a many-to-many relationship: a gene may link to multiple enhancers and an enhancer may link to multiple genes. We tested three types of epigenomic assays to approximate *A*_*r*_ by counting sequencing reads in the enhancer: DNase-seq, ATAC-seq, or H3K27ac ChIP-seq. The contact *C*_*r,g*_ is taken from a normalised chromatin contact frequency matrix. A pseudocount is added to each *C*_*r,g*_, so that all candidates *R*_*g*_ are taken into account (see Supplementary Material).

### 2.2 Generalised Activity-By-Contact-score

We present a two-fold adaptation of the ABC-score, to account for all candidate target genes of an enhancer and for all transcription start sites (TSSs) of a gene. Regarding the former, the activity of an enhancer conceptually represents all regulatory interactions an enhancer has. However, an enhancer can interact with different genes and not all genes are equally likely to be brought into vicinity of the enhancer. We assume that an enhancer’s regulatory input is a function of the number of contacts with all potential target genes. Target genes that are often in contact with the enhancer would receive more regulatory input than genes with fewer contacts. Thus, the activity *A*_*r*_ could be denoted in a relative manner:

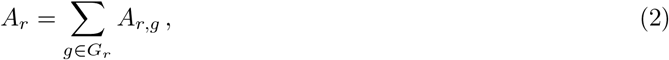

where *A*_*r,g*_ denotes the relative regulatory activity of enhancer *r* towards gene *g* and *G*_*r*_ denotes the set of all genes that are located within a predefined window around *r*. Since the values of *A*_*r,g*_ are not known, we propose an approximation by using chromatin contacts:

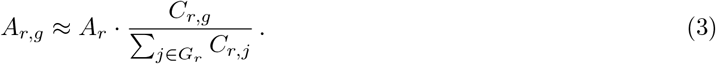

We approximate the regulatory activity *A*_*r,g*_ by the relative fraction of its contact *C*_*r,g*_ to the TSS of *g* to the sum over the contacts to all genes *G*_*r*_ in a window around the enhancer. Thus, the ABC-score becomes:

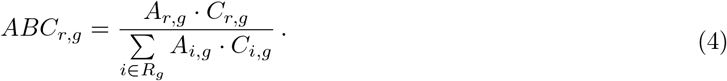

In comparison to Eq. (1), the activity *A*_*r*_ of an enhancer is replaced with its gene-specific relative activity *A*_*r,g*_, see Eq. (3). Thus, this adapted score does not only function in a gene-centric way, but also in an enhancer-centric way, by adapting the activity to the relative number of contacts of an enhancer’s candidate target genes. This adaptation therefore uses a reduced enhancer activity estimate, in particular for enhancers in contact with many genes, and prevents them from being accounted with their full activity for all genes within the window.

In addition, instead of considering only one TSS per gene, e.g. the 5’ TSS, we propose to include all TSSs of a gene in the following fashion:

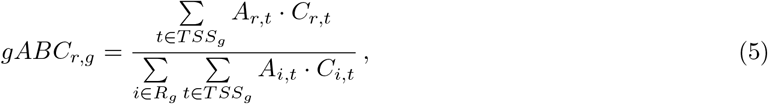

where *TSS*_*g*_ are all annotated TSSs of gene *g*. That allows to include the contact information to more potentially relevant transcription sites and omits the selection of an individual TSS. We name the score with these two changes the generalised ABC (gABC) score.

### 2.3 Validation on CRISPR-screens

To validate our generalised ABC-score and to test different assays for measuring enhancer activity, we examined the performance on experimentally validated enhancer-gene interactions. We made use of three CRISPRi-screens for K562 cells. Gasperini et al. (2019) used a single-cell CRISPRi-screen, introducing guide RNAs at a high multiplicity of infection, followed by single-cell RNA-seq. Schraivogel et al. (2020) developed targeted Perturb-seq (TAP-seq), which promises to be more sensitive by targeting genes of interest for the transcriptomic readout, and was established by a screening on two chromosomes. Fulco et al. (2019) used their CRISPRi-FlowFISH approach and collected data from other CRISPR-based studies. Unlike Fulco et al. (2019), we neither divide interactions into enhancer-gene and promoter-gene pairs nor exclude interactions, where the expression decreased after enhancer perturbation. We tested four different set-ups for measuring enhancer activity: 1) DNase-seq, 2) H3K27ac ChIP-seq, 3) the geometric mean of DNase-seq and H3K27ac ChIP-seq, 4) and ATAC-seq. In addition, we assessed a K562 Hi-C matrix, a Hi-C matrix averaged across 10 cell types constructed by Fulco et al. (2019), and a contact estimate (inverse of the linear distance) based on a fractal globule model (Lieberman-Aiden et al., 2009). For each combination we evaluated the ABC- and gABC-scoring approach and calculated precision-recall curves. In addition, we tested the significance of the pairwise difference between the area under the receiver operate characteristic (ROC) curves (DeLong et al., 1988, Robin et al., 2011) (more details in Supplementary Material).

We also compared gABC to Enformer, a sequence-based deep learning model predicting gene expression and chromatin states, whose characteristic is an increased information flow between distal sequence positions (Avsec et al., 2021). To quantify enhancer-gene interactions with Enformer we compared the expression estimate upon *in silico* mutagenesis of the enhancer region, by either replacing 2 kb centred at the enhancer with neutral nucleotides, or by shuffling the sequence for 25 iterations (Karollus et al., 2022). Due to the size of Enformer’s receptive field, we limited the comparison to interactions with ≤96 kb distance.

### 2.4 TF affinities and summarisation on gene-level

In addition to enhancer-gene interactions, STARE aims to describe a TF’s regulatory impact on a gene (Fig. S1). There are two steps required. First, genomic regions that influence the regulation of a gene have to be identified. This can be done via ABC-scoring as described before, or in a more simplistic approach by taking all open regions within a defined window around the gene’s TSS into account. Utilising the ABC-model allows STARE to function with any desired window. Second, the affinities of TFs to the identified regions have to be quantified. Instead of relying on calling TF binding sites, we use the tool TRAP, which calculates relative binding affinities for TFs in a genomic region. TRAP describes TF binding with a biophysical model to predict the number of TF molecules that bind in a sequence (Roider et al., 2007). The higher the affinity, the more likely a TF is to bind. Retaining low binding affinities can hold valuable information (Schmidt et al., 2016, Kribelbauer et al., 2019) and omits selection of an arbitrary threshold. For all analyses presented in this manuscript we used a non-redundant collection of 818 human TF motifs in the form of position frequency matrices (PFMs) from JASPAR 2022, HOCOMOCO v11, and the work of Kheradpour and Kellis (Castro-Mondragon et al., 2022; Kulakovskiy et al., 2018; Kheradpour and Kellis, 2014). When converting PFMs to position-specific energy matrices required by TRAP, we take the average nucleotide content of the candidate regulatory regions as background.

To summarise the TF affinities in enhancers per gene, we combine them with the predicted enhancer-gene interactions. The summarisation depends on how the region-gene mapping was done. For the window-based approach, TF affinities in all open regions around the gene’s TSS are summed:

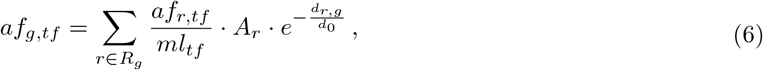

where *af*_*g,tf*_ is the affinity of TF *tf* summarised for gene *g. R*_*g*_ is the set of all open regions *r* that were located within the window around *g. af*_*r,tf*_ is the affinity of *tf* in *r, ml*_*tf*_ is the motif length of *tf* and *A*_*r*_ is the activity for *r*. The affinity is corrected for the distance *d*_*r,g*_ of *r* to the TSS of *g* by an exponential decay function, as proposed by Ouyang et al. (2009), where *d*_0_ is set to a constant of 5000 bp.

When the gABC-score was used to assign regions to genes, the summarisation changes, as there is more epigenomic information available. Regions close to the TSS (≤2500 bp) are always included, independently of their gABC-score, as they are very informative for the expression regulation of a gene (Schmidt et al., 2019). They are also scaled differently:

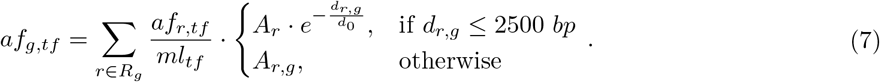

*R*_*g*_ is the set of regions that was linked to the gene with the gABC-score and *A*_*r,g*_ is the adapted activity (Eq. 3). For regions close to the TSS the base activity *A*_*r*_ is corrected with the exponential decay function, as the contact frequency would likely be the contact of the region with itself and thus *A*_*r,g*_ could be erroneous. When using the regular ABC-score, the affinity scaling changes as follows:

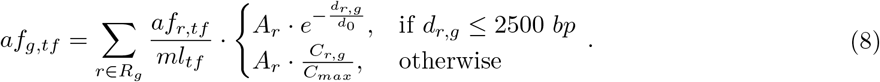

The regions close to the TSS are scaled in the same way as before. For all the other regions we divide the contact frequency *C*_*r,g*_ by the maximum contact that was measured for all region-gene pairs *C*_*max*_. The reasoning is to incorporate the contact frequency and to have both multipliers for *A*_*r*_ in the range of [0, 1]. Essentially, the activity multipliers for gABC and ABC differ only in how the contact is scaled: for gABC relative to all gene contacts of the respective enhancer and for ABC to the contacts of all enhancer-gene pairs. In addition to the TF affinities, we report three additional gene features: the number of regions considered per gene, the regions’ average distance to the TSS, and the regions’ average base pair length, as all three can be predictive of gene expression (Schmidt and Schulz, 2019).

### 2.5 Application to single-cell data

To test the capability of gABC-scored enhancer-gene interactions combined with TF affinities to capture regulatory cell type-specific information, we analysed a single-cell data set of the human heart from Hocker et al. (2021), providing single-nuclei (sn) ATAC-seq, as well as snRNA-seq data. The candidate enhancers were pooled and ATAC-seq RPKM was measured for each cell type. For chromatin contacts we tested H3K27ac HiChIP data of the left ventricle (Anene-Nzelu et al., 2020), as well as an average Hi-C matrix (Fulco et al., 2019). Regions known to accumulate an anomalous amount of sequencing reads were excluded (Amemiya et al., 2019, The ENCODE Project Consortium, 2012). As we had enhancer activity and enhancer contact data at hand, we determined enhancer-gene interactions for each cell type with the gABC-score (cut-off 0.02, 5 MB window around the 5’ TSS), and summarised TF affinities for each gene in each cell type based on those interactions. To assess the predictability of gene expression by gene-TF affinities, we used INVOKE (Schmidt et al., 2016), which implements a linear regression model based on gene-TF scores, and selects TFs that are most predictive of gene expression. We trained prediction models for multiple set-ups for each cell type to compare their prediction accuracy. We repeated this process for cell type-specific (CS) genes, defined as genes where the z-score of expression between cell types was ≥2 and TPM ≥0.5. The gene-TF matrices were limited to TFs expressed in a cell type (TPM ≥ 0.5). Schmidt et al. (2020) applied INVOKE on similar data types for bulk data, but restricted information of distant enhancers to those connected to promoters via loops. With the ABC-approach we have a finer resolution of regulatory interactions and can integrate the contact frequency into the summarisation of TF affinities.

#### 2.5.1 Comparison to co-accessibility analysis

A common approach to identify regulatory interactions in single-cell ATAC-seq data is to call co-accessible regions. Hocker et al. (2021) ran Cicero (Pliner et al., 2018) on their snATAC-seq data to derive pairs of regions with correlated accessibility, limited to a distance of 250 kb. Whenever either side of a co-accessible region pair overlapped a 400 bp window around any annotated TSS of a gene, we considered it as an enhancer-gene interaction for that gene. We tested how informative the resulting 62,384 co-accessible interactions are for our gene expression prediction model. Summarisation of TF affinities was done according to Eq. 6, and the affinities of regions close to the TSS (≤ 2500 bp) were included, as described in Sec. 2.4.

#### 2.5.2 Intersection with eQTLs

Further, we compared the agreement of the regular ABC- and gABC-score with eQTL data. We intersected ABC-scored interactions from four different heart cell types (Hocker et al., 2021) and K562 cells with eQTL-gene pairs of matching samples from the GTEx portal (The GTEx Consortium, 2020). We used high confidence eQTL-gene pairs from three different fine-mapping approaches, namely CAVIAR, CaVEMaN and DAP-G (Hormozdiari et al., 2014, Brown et al., 2017, Wen et al., 2016). For each set of eQTLs, we defined the enhancer-gene pairs that were supported by the eQTLs, meaning all candidate enhancers of a cell type with a variant, where the affected target gene was within the chosen ABC window size. Then, we compared the fraction of those eQTL-supported enhancer-gene pairs that we could also find among a variable number of highest scored ABC/gABC interactions (Recall). As we also had the co-accessibility analysis on the heart data, we examined how many eQTL-gene pairs the resulting interactions recover.

## 3 Results

### 3.1 Generalised ABC-score improves interaction prediction

We propose a generalised ABC-score (Eq. 5), where the activity of an enhancer is described in a gene-specific manner (Fig. 1c, Eq. 3), and all annotated TSSs of a gene are considered. On all validation data sets and for all combinations of activity measurements the gABC-score outperformed the regular ABC-score (p-value ≈0.0005 Wilcoxon signed-rank test) (Fig. 1a+d; Tab. 1). The difference was more pronounced in the Gasperini and Schraivogel validation data. Each of the two adaptations of the gABC-score individually increased the area under the precision recall curve (AUPRC) compared to the regular ABC across activity assays, with the gene-specific activity providing an average gain of 0.026, and including all TSSs giving an average improvement of 0.053. Taken together, the gABC-score yielded on average a 0.107 higher AUPRC (Tab. S1). The areas under the ROC curves for gABC were significantly higher in ten out of twelve pairwise comparisons across CRISPRi-screens and activity assays (Tab. S5). Using an average Hi-C matrix changed the accuracy marginally for both ABC-scores (Fig. S2a; Tab. 1). When using the fractal globule module to estimate contact frequency only based on distance, gABC achieved less improvement over the regular ABC with an average AUPRC gain of 0.048 (Tab. S1). We could reproduce the higher accuracy of the gABC-score in a direct comparison to the implementation of Fulco et al. (2019) (Tab. S2). Further, we examined the correlation between each of the ABC-scoring approaches and the absolute change in gene expression as measured in the CRISPRi-screens. The gABC-score showed a higher Spearman correlation coefficient across all three data sets (Tab. S3).

**Figure 1.**
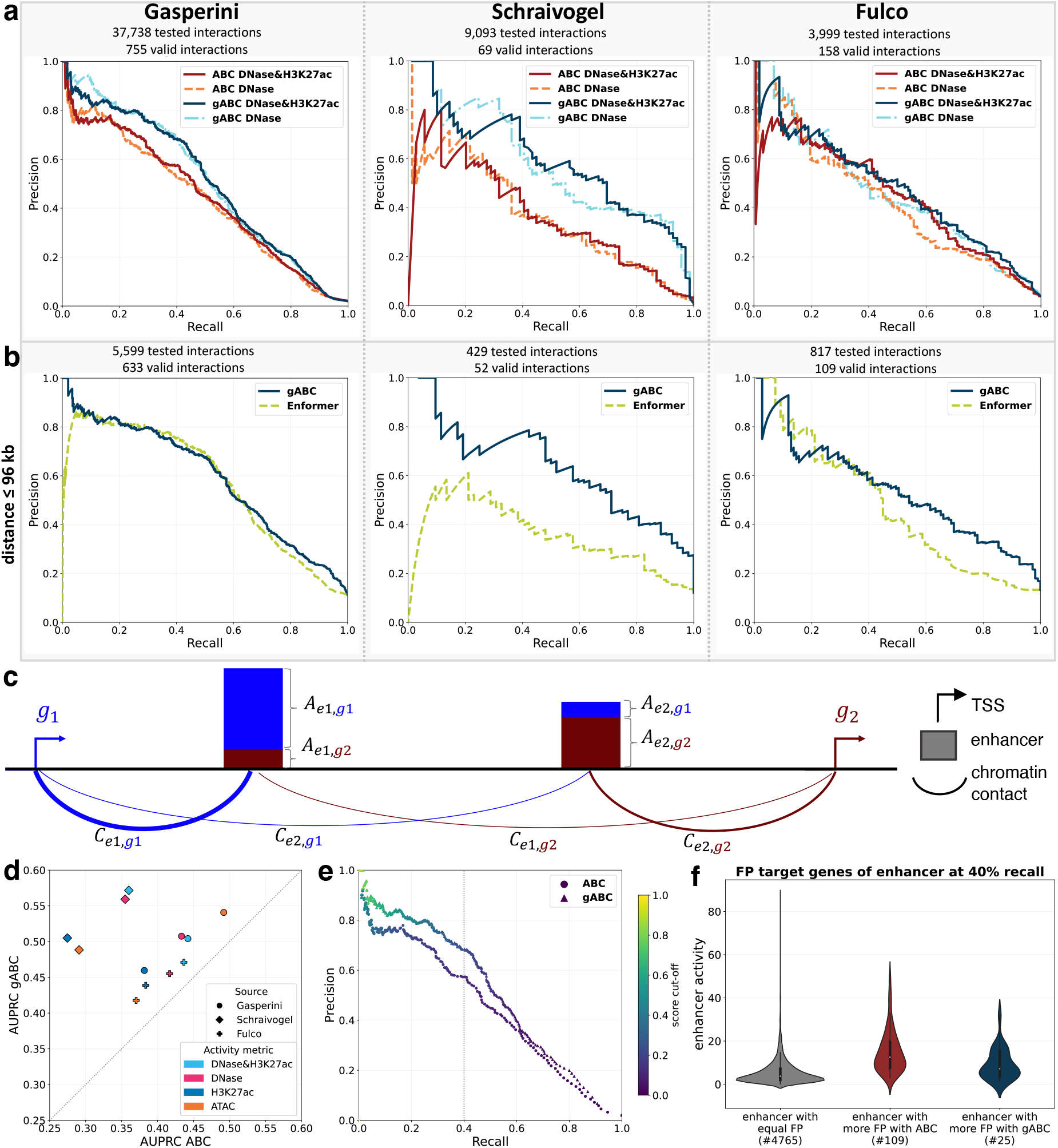
Performance comparison of the ABC- and gABC-score on CRISPRi-screens in K562 cells using different epigenomic assays. For chromatin contacts a K562 Hi-C matrix (5 kb resolution) was used (Rao et al., 2014). **(a)** Precision-recall (PR) curves of both ABC-scores on experimentally validated enhancer-gene links. The AUPRC values can be found in Tab. 1. **(b)** PR curves comparing gABC with Enformer on interactions with a distance of ≤96 kb. Out of four tested calculations for the predicted expression difference of Enformer the best one is shown. The AUPRC values are listed in Tab. S4. DNase-seq and H3K27ac ChIP-seq was used as activity for gABC. **(c)** Schema for the gene-specific enhancer activity, which distributes the activity of an enhancer among its scored genes, dependent on contact frequencies. **(d)** Direct comparison of the AUPRC for ABC and gABC on different CRISPRi-screens and with different assays for enhancer activity. **(e)** PR curve coloured by ABC/gABC score respectively with DNase&H3K27ac as activity on the CRISPRi-screen of Gasperini et al. (2019). The dotted grey line marks the position at 40% recall. **(f)** Distribution of the activity of enhancers (geometric mean of read counts of DNase-seq and H3K27ac ChIP-seq) separated by the number of false positive (FP) target genes called by each method at 40% recall on the screen of Gasperini et al. (2019). ‘equal FP’ contains all enhancers where the number of FP target genes is the same for both scores (0 FP included). ‘more FP’ means that either of the scores called more FP target genes for that enhancer. The number of enhancers in each category is shown below the x-axis labels.

**Table 1.** AUPRC using ABC and gABC for identifying regulatory interactions on three validation data sets, with different assays for enhancer activity and contact data. The highest AUPRC within each column is written in bold.

| Validation data | Gasperini et al. (2019)<br>755 valid / 37,738 interactions |  |  |  | Schraivogel et al. (2020)<br>69 valid / 9,093 interactions |  |  |  | Fulco et al. (2019)<br>158 valid / 3,999 interactions |  |  |  |
| --- | --- | --- | --- | --- | --- | --- | --- | --- | --- | --- | --- | --- |
| Enhancer activity assay | DNase& H3K27ac | DNase | H3K27ac | ATAC | DNase& H3K27ac | DNase | H3K27ac | ATAC | DNase& H3K27ac | DNase | H3K27ac | ATAC |
| ABC K562 Hi-C | 0.4421 | 0.4333 | 0.3816 | 0.4919 | 0.3600 | 0.3549 | 0.2746 | 0.2909 | 0.4365 | 0.4167 | 0.3835 | 0.3702 |
| gABC K562 Hi-C | **0.5042** | **0.5076** | 0.4596 | **0.5408** | **0.5717** | **0.5592** | **0.505** | **0.4884** | **0.471** | **0.4552** | **0.4388** | **0.4176** |
| ABC avg Hi-C | 0.4486 | 0.4395 | 0.3898 | 0.4929 | 0.3554 | 0.3661 | 0.2806 | 0.2945 | 0.436 | 0.4085 | 0.386 | 0.3663 |
| gABC avg Hi-C | 0.5038 | 0.5072 | **0.4605** | 0.5364 | 0.5552 | 0.5507 | 0.4985 | 0.475 | 0.452 | 0.4387 | 0.4259 | 0.4015 |

To disentangle for which enhancers the gABC-score performs better than the regular ABC, we focused on the largest CRISPRi-screen from Gasperini et al. (2019) (Fig. S3a) and found that the regular ABC predicted more false positive target genes for enhancers with a high activity (Fig. 1e+f, Fig. S3b-e). There was a small subset of enhancers in gene-rich regions for which the gABC-score predicted more false positive interactions (Fig. S3f-i).

Enformer is a novel method that predicts gene expression and chromatin states directly from the DNA sequence using a complex neural network architecture outperforming other sequence-based models (Avsec et al., 2021). We compared the gABC-score and an Enformer model learned on K562 cells using *in silico* mutagenesis, where the strength of interactions was quantified by the predicted expression change upon sequence perturbation of the enhancer. gABC achieved a higher accuracy on all three validation data sets (Fig. 1b) testing alternative ways for the *in silico* mutagenesis (Tab. S4). This was also reflected in significantly higher areas under the ROC curves (all p-values ≤ 0.05, DeLong et al., 1988, Robin et al., 2011).

Although it uses the same information, the gABC-score performed better than the regular ABC-score and outperformed the accurate sequence-based Enformer model.

### 3.2 One activity assay yields similar accuracy

The original formulation of the ABC-score requires two types of assays to measure enhancer activity, namely DNase-seq and H3K27ac ChIP-seq (Fulco et al., 2019). We tested the performance of the gABC-scoring principle using different assays for measuring enhancer activity (Fig. 1a+d; Tab. 1; Fig. S2b). On the data sets of Schraivogel and Fulco the combination of DNase-seq and H3K27ac ChIP-seq performed better than either of them alone, although the performance drop with only DNase-seq was small. On the Gasperini data, DNase-seq without H3K27ac ChIP-seq was slightly better. Quantifying enhancer activity with ATAC-seq resulted in a worse performance on the Schraivogel and Fulco validation data, but a clear improvement on the Gasperini data set, surpassing the combination of DNase-seq and H3K27ac ChIP-seq.

### 3.3 Regulatory interactions in single-cell data

Using just one epigenome assay for the ABC-score enables direct application on high-resolution snATAC-seq data, which has the potential to improve the prediction of cell type-specific interactions. As a proof of concept, we analysed a human heart snATAC-seq data set, comprising eight cell type clusters (Hocker et al., 2021). The candidate enhancers of the cell types were pooled to a set of 286,777 regions with a summarised ATAC-seq measurement for each defined cell type (Fig. 2a). On average we predicted 408,846 gABC-scored interactions (SD≈ 12,200) over all cell types. Approximately 23.6% of a cell type’s interactions were shared with all other cell types (Fig. 2b; Fig. S4a). Each cell type also featured unique interactions although this was highly variable (*μ*≈ 11.6%, SD≈5.1%). Atrial cardiomyocytes (aCM) and ventricular cardiomyocytes (vCM) formed the largest intersection of interactions found in only two cell types, consisting of 39,707 interactions. The average median of enhancers per expressed gene (TPM ≥ 0.5) across cell types was 4.75 (SD≈ 0.43) (Fig. S4b). Despite all cell types having the same candidate enhancers and a shared contact measurement, their predicted enhancer-gene interactions appeared to be considerably distinct.

**Figure 2.**
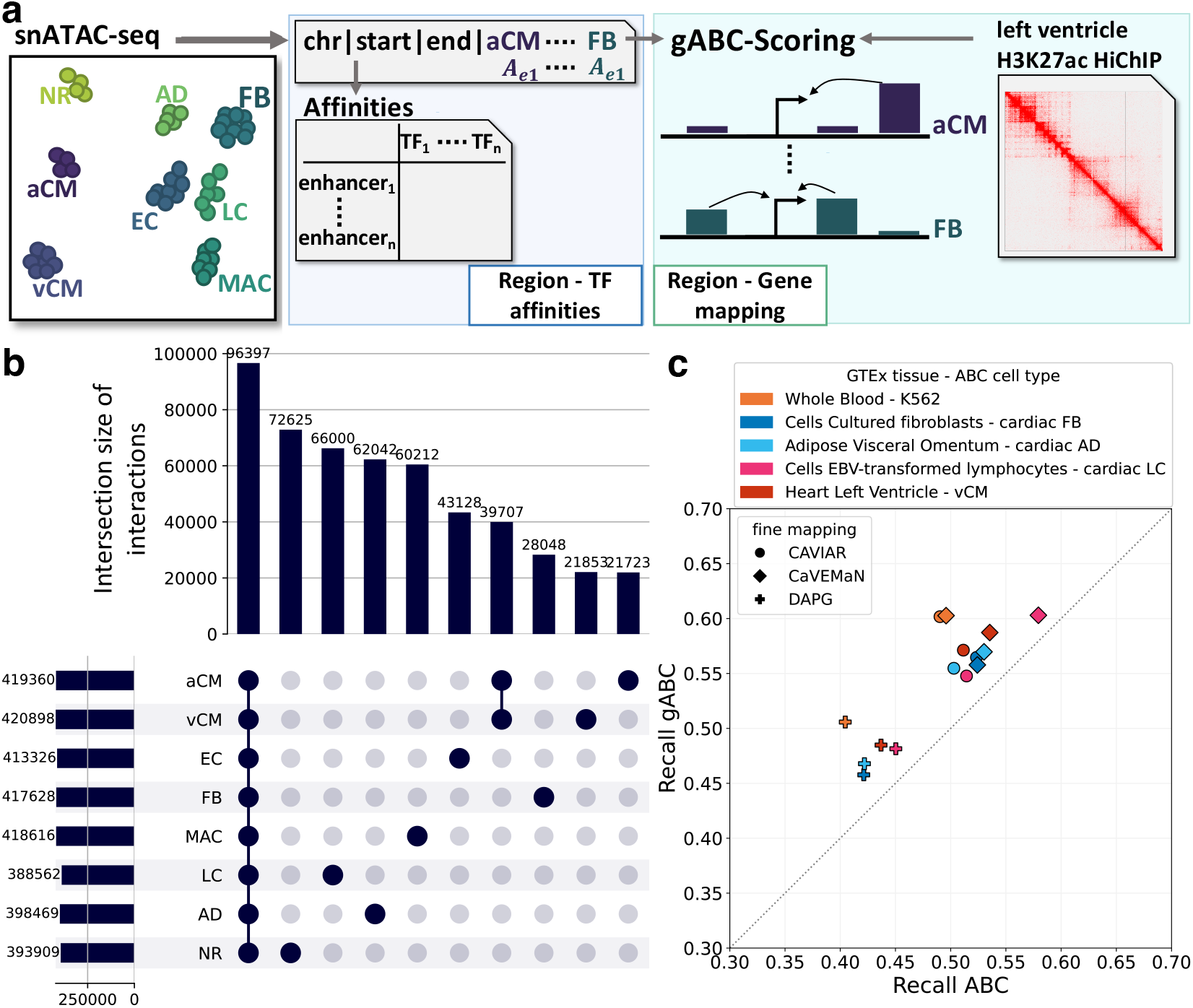
Enhancer-gene interactions called in single-cell heart data. **(a)** Schema of data processing. snATAC-seq from Hocker et al. (2021) was used to identify candidate enhancers and their activity in the annotated cell types. For each cell type gABC-interactions were called. Enhancer-gene contacts were retrieved from left ventricle H3K27ac HiChIP data (Anene-Nzelu et al., 2020). **(b)** Upset plot of the enhancer-gene interactions called in each cell type. Only the 10 largest intersections are shown. **aCM**: atrial cardiomyocytes; **vCM**: ventricular cardiomyocytes; **EC**: endothelial cells; **FB**: fibroblasts; **MAC**: macrophages; **LC**: lymphocytes; **AD**: adipocytes; **NR**: nervous cells. **(c)** Intersection of eQTL-gene pairs from different GTEx samples with the ABC- and gABC-interactions. Recall is the fraction of enhancer-gene pairs found by each score out of all pairs where the enhancer contained an eQTL whose target gene was within the window size used for ABC-scoring. The 300,000 highest scored interactions of ABC and gABC were used.

#### 3.3.1 gABC-score recovers more eQTL-gene pairs

We examined how many eQTLs from different tissues are recovered by ABC interactions from matching cell types, including four heart cell types from Hocker et al. (2021) and K562 cells. We took high confidence eQTL-gene pairs from three different fine-mapping methods and compared which fraction of enhancer-gene interactions supported by eQTLs were also found by the 300,000 highest scored ABC- and gABC-interactions (Fig. 2c). The gABC-interactions recovered significantly more eQTL-gene pairs across all fine-mapping methods and eQTL data sets (p-value≈0.0005 Wilcoxon signed-rank test). This finding was reproduced for the 100,000 and 200,000 highest scored interactions. The gABC-interactions also captured more eQTL-gene pairs than interactions derived via co-accessibility analysis (Fig. S4c).

#### 3.3.2 Linking epigenetic features to cell type-specific expression

We trained cell type-specific gene expression prediction models based on gene-TF affinity matrices, constructed with different approaches (Fig. 3a). Using the gABC-score performed best across all cell types (Pearson correlation coefficient *μ*≈0.563), with similar results using the average Hi-C matrix (*μ*≈0.556). The regular ABC-score had a slightly lower performance (*μ*≈0.538), whereas interactions identified via co-accessibility resulted in the lowest performance (*μ*≈0.505). Since the co-accessible interactions were limited to a distance of 250 kb, we also tested the gABC-score in a 500 kb window, which marginally decreased the performance in the prediction model (*μ*≈0.562, Fig. S5a).

**Figure 3.**
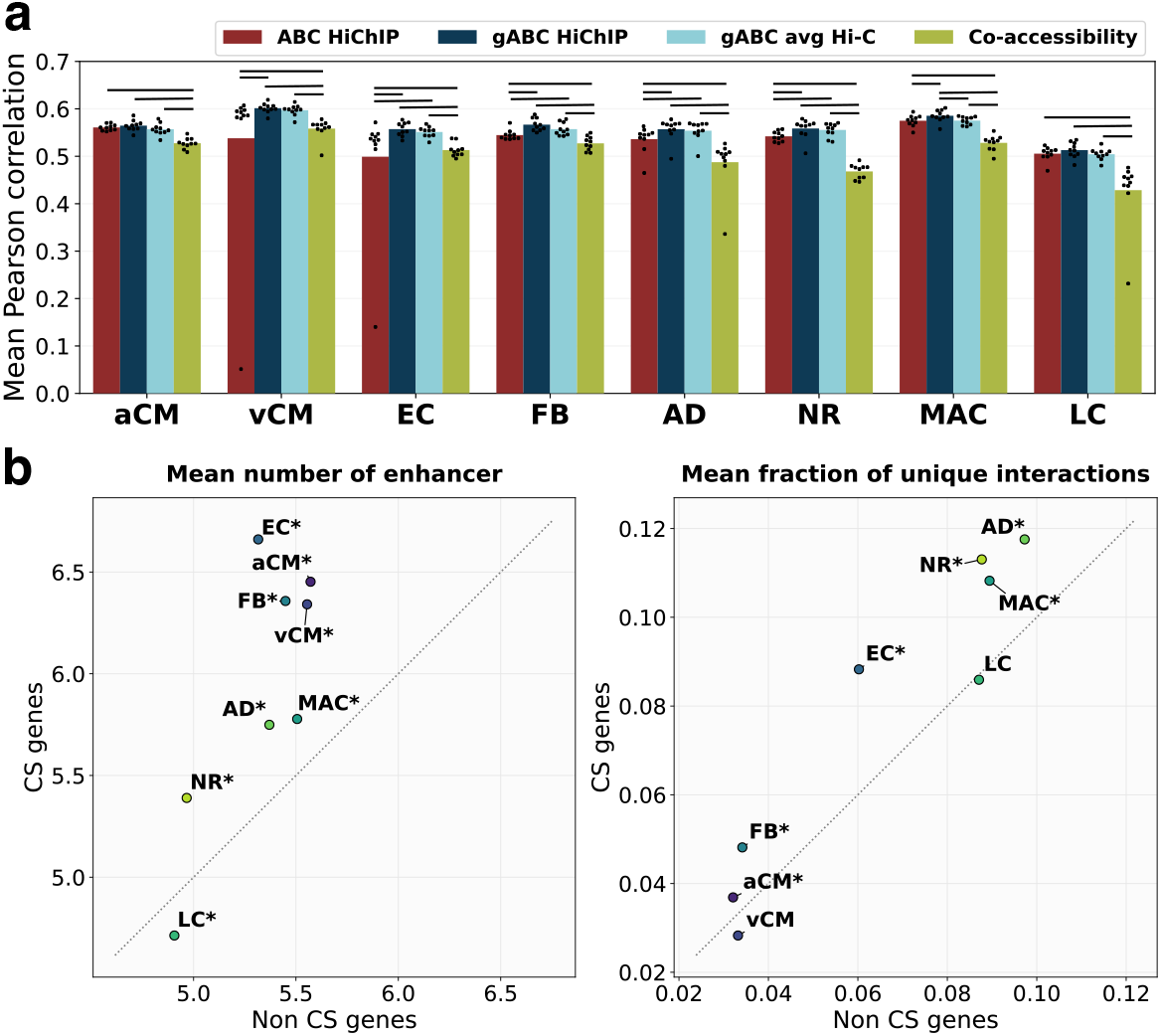
Gene expression prediction on single-cell data and characterisation of CS genes. **(a)** Accuracy of a gene expression prediction model based on different gene-TF affinity matrices. The model was trained on all genes with available expression values. The Pearson correlation coefficient is shown as average over a 10-fold outer cross-validation. A horizontal line above the bars indicate significance (p-value≤ 0.05, Mann-Whitney U test). snATAC-seq data was used as enhancer activity for all approaches. ABC/gABC H3K27ac HiChIP: regular/generalised ABC scoring with H3K27ac HiChIP as contact data; gABC avg Hi-C: gABC with an average Hi-C matrix as contact data; Co-accessibility: enhancer-gene links defined by Cicero (Pliner et al., 2018), see Sec. 2.5.1. The respective Spearman correlation coefficients, as well as additional approaches and their performance in a training on CS genes only, are presented in Fig. S5. **(b)** Comparison of attributes between CS (TPM ≥ 0.5 and z-score ≥ 2) and not cell type-specific (non-CS) genes (TPM ≥ 0.5) (*: p-value ≤ 0.05, Mann-Whitney U test). See Fig. 2 for cell type abbreviations.

We defined a set of CS genes (TPM≥0.5 and z-score≥2) for each cell type (*μ*=1,404 genes, SD≈582 genes) and analysed those in more detail. The CS genes were mostly unique to a cell type (Fig. S4d), and GO term enrichment returned cell-type appropriate terms (Fig. S4e). To further characterise the sets of CS genes, we examined additional attributes and compared CS genes to non-CS genes, both sets restricted to expressed genes (TPM≥0.5) (Fig. 3b; Fig. S4f). CS genes had more assigned enhancers than non-CS genes in all cell types except for LC. This matches the finding of Fulco et al. (2019), who described a higher number of enhancers for tissue-specific than for ubiquitously expressed genes. Furthermore, CS genes tended to have a higher percentage of unique interactions, meaning the fraction of interactions of a gene, that were exclusively found in that cell type, was higher. Most cell types had a slightly higher average activity in enhancers linked to CS genes than in enhancers linked to non-CS genes, except for aCM and FB. There was no clear trend visible for the average contact frequency or TSS distance of the assigned enhancers. To exclude that the differences were solely caused by the CS genes’ higher expression, we repeated the comparisons but restricted them to CS and non-CS genes from the upper quartile of TPM values. The results were highly similar across all features (data not shown). We repeated the training of gene expression models but limited it to CS genes and observed an overall decrease in prediction accuracy (Fig. S5b). The gABC-score still performed better than the ABC-score. The prediction model returns a regression coefficient for each TF, indicative of TF relevance, and allows to investigate which TFs might drive gene expression in which cell type (Fig. S5c). Overall, we were able to characterise cell type-specific expression regulation based on single-cell chromatin accessibility and bulk chromatin contact data.

### 3.4 Runtime of STARE

We optimised STARE’s runtime by allowing multiple steps to be run in parallel (Fig. 4a) and to omit redundant calculations, if multiple cell types/metacells/individual cells with a respective activity measurement are processed. The runtime per activity column decreases when multiple columns are handled in the same run, allowing large data sets to be processed in a few minutes (Fig. 4b).

**Figure 4.**
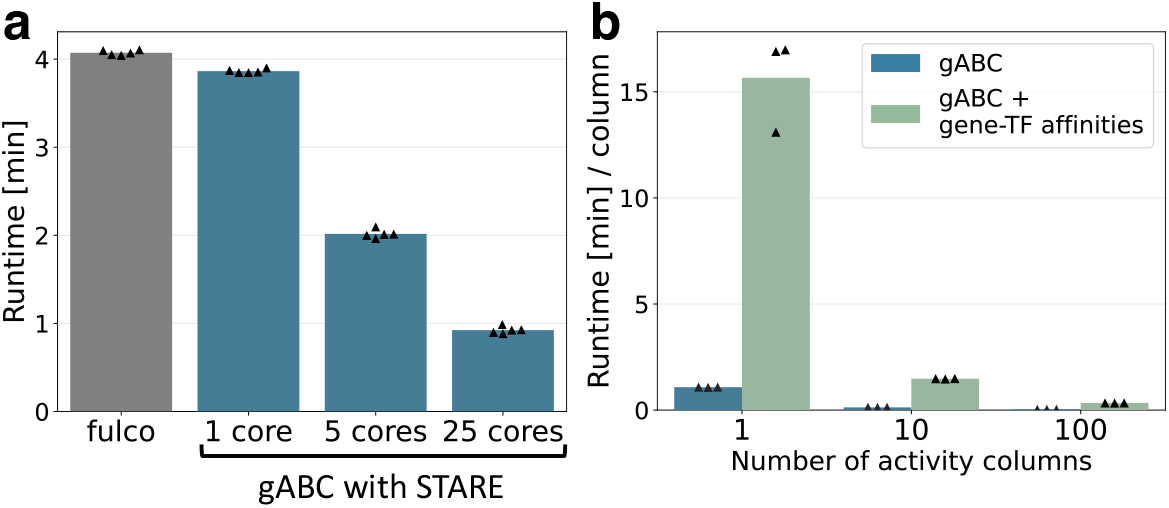
Runtime of the STARE pipeline. **(a)** Runtime of the original ABC pipeline by (Fulco et al.) and our ABC-scoring implementation with a different number of cores. Any writing of output files was omitted. Calculations were done for K562 cells, scoring 155,976 candidate enhancers for 24,586 genes in a 10 MB window. The bars show the mean of five runs. **(b)** Runtime divided by the number of activity columns for the ABC-scoring alone and when additionally calculating the gene-TF affinity matrix. Writing and compression of output files is included. Calculations were done on single-cell heart data (Hocker et al., 2021) with 286,777 candidate enhancers, for 55,765 genes in a 5 MB window. 818 TFs were assessed. The bars’ height represent the mean of three runs.

## 4 Discussion

We present a variation of identifying regulatory enhancer-gene interactions with the ABC-model. In its original study the ABC-model already showed a better accuracy for detecting validated enhancer-gene links than other approaches (Fulco et al., 2019). We propose a generalised ABC-score where the activity of an enhancer is described in a gene-specific manner by weighting it relative to the number of enhancer-gene contacts and where all TSSs of a gene are considered. This generalisation resulted in an improvement in identifying experimentally validated enhancer-gene interactions, both compared to the regular ABC, as well as to the deep learning model Enformer. It should be noted however, that Enformer was built to predict gene expression, and not to identify enhancer-gene links (Avsec et al., 2021). The accuracy of different scoring approaches was partially inconsistent between validation data sets, especially when using ATAC-seq for enhancer activity. Although all data sets are based on a dCas9-KRAB system, there are differences in the experimental set-ups and scope. Gasperini et al. (2019) introduced multiple guide RNAs and measured expression via single-cell RNA-seq, allowing quantification of interactions for over 10,000 genes. Schraivogel et al. (2020) presented their method TAP-seq, while Fulco et al. (2019) published CRISPRi-FlowFISH and collected previous results, both evaluating interactions for less than one hundred genes. Details in candidate enhancer selection, filtering steps, and processing might lead to biases or varying sensitivity in the identification of enhancer-gene pairs, which is already indicated by the different fractions of significant interactions. Further, it is debatable whether CRISPRi-screens are able to detect all regulatory interactions, as true enhancers with small effect sizes may be overlooked. Moreover, perturbations of individual enhancers are presumably not capable of accounting for shadow enhancers with redundant functionality (Schoenfelder and Fraser, 2019; Singh and Yi, 2021). More large-scale validation data could consolidate and contextualise our findings, but are currently unavailable. Importantly, any model to annotate enhancer-gene interactions is only a prediction and likely not capturing the whole regulatory complexity of genes. The ABC-model requires two data types, which makes it applicable in a range of scenarios, but it might also miss out relevant epigenetic information. Further, the model assumes all genes are regulated in the same way.

In context of required data types, we were able to demonstrate that, unlike suggested in the original work, one assay for measuring enhancer activity works similarly well, specifically using DNase- or ATAC-seq without H3K27ac ChIP-seq data. Further, using averaged contact data yielded a high performance as well, which broadens the applicability of the ABC-score to all data sets, where a measurement of enhancer activity is available. Especially for single-cell epigenomics it is challenging to measure multiple modalities in the same cell.

We present the STARE framework to derive gene-TF affinities. After mapping candidate enhancers to genes, using either the ABC-score or a window-based approach, STARE summarises TF affinities on a gene level. Unlike other methods aiming to determine regulatory relations of TFs to genes (Lan et al., 2012; Wang et al., 2013), STARE does not require scarce TF ChIP-seq data. It uses a motif-based biophysical model (Roider et al., 2007) to determine TF affinities in accessible regions. As consequence, unlike other methods (McLeay et al., 2012), it is able to retain low affinity binding information. Patel and Bush (2021) use similar data as STARE and rely on a graph-based approach, but they do not incorporate active regulatory regions and analysis is limited to a window marked by the most distant CTCF peaks within 50 kb of the gene body. With the ABC-model, STARE specifically determines candidate enhancers within any selected window size. Notably, our model assumes an additive influence of TFs, which is not likely to accurately capture biological reality (Zeitlinger, 2020). Furthermore, there is potential redundancy on two levels: the aforementioned functional redundancy of enhancers and the redundancy of TF binding motifs (Gitter et al., 2009; Cusanovich et al., 2014).

We applied our framework to single-cell data with clustered cell types of the human heart (Hocker et al., 2021), where the activity of a unified set of candidate enhancers was measured for each cell type. The assumption was that cell type-specificity is mainly driven by activity of regulatory regions and less by spatial chromatin conformation. Although chromatin contacts were also found to change upon cell differentiation (Fraser et al., 2009; Zhang et al., 2020), the 3D conformation of the genome is described as less dynamic and more as a scaffold to enable and stabilise regulatory interactions (Schoenfelder and Fraser, 2019; Ing-Simmons et al., 2021). We were able to unravel differences in regulatory interactions across cell types and to characterise regulation of cell type-specific genes, despite using bulk chromatin contact data. We demonstrated a downstream application example of STARE with a linear expression model, which allows to identify candidate regulatory TFs for further evaluation. Considering the small training sets and that the model assumes the same regulation for all genes, the prediction yielded a reasonable performance. Schmidt et al. (2020) used a similar expression prediction approach and incorporated distant enhancers. However, their work relies on annotated loops, which are not likely to cover all relevant regulatory interactions. In addition, their strategy is not able to integrate contact frequencies into TF affinities, nor to derive cell type-specific interactions from bulk contact data. There are other tools for predicting gene expression explicitly in individual cells that incorporate TF information, such as SCENIC (Aibar et al., 2017), ACTION (Mohammadi et al., 2018) or TRIANGULATE (Behjati Ardakani et al., 2020), but none of them considers long-range enhancer-gene interactions. A compelling approach would be to combine these expression prediction tools with information on distant enhancers.

STARE represents a currently unique form of deriving TF affinities on a gene level: it combines enhancer-gene links called by the ABC-score with a non hit-based TF annotation. Prospectively, we would like to apply STARE on individual cells instead of clustered cell types, which would require additional steps to account for the drastic sparsity of most single-cell measurements. It would also be highly interesting to further investigate the importance of chromatin contacts in cell type-specificity, once the resolution and availability for such data are advanced.

## Availability and genome version

The presented results are provided via Zenodo (https://doi.org/10.5281/zenodo.5841991). All data is in hg19. For details on the used data see Supplementary Material.

## Acknowledgements

We thank Nina Baumgarten for advice and discussions on the implementation. We acknowledge the ENCODE Consortium and the lab of Michael Snyder in Stanford who generated the ATAC-seq data sets.

