## Supplementary Material for "The adapted Activity-By-Contact model for enhancer-gene assignment and its application to single-cell data"

#### 1 Validation on CRISPRi-screens

We examined the performance of different ABC-scoring setups on three experimentally validated enhancer-gene interactions in K562 cells from Gasperini et al. (2019), Schraivogel et al. (2020) and Fulco et al. (2019). From all studies we took interactions with a false discovery rate (FDR) of $\leq 5\%$ as our validation data set. For the data from Schraivogel et al. (2020) we first filtered for interactions with a distance of $\leq 5$ MB and kept interactions to genes with at least one significant associated enhancer, similar to the original publication. To have the same set of candidate enhancers for all CRISPRi-screens, we obtained the enhancers from Fulco et al. (2019) who defined them as follows: each DNaseI hypersensitive site was extended to a total length of 500 bp and in addition, all promoter regions of 500 bp around the transcription start site (TSS) of all genes were included. Only interactions that intersected at least one K562 candidate enhancer could be considered. Regions known to accumulate an anomalous amount of sequencing reads were excluded (Amemiya et al., 2019, The ENCODE Project Consortium, 2012). In addition to DNase-seq, H3K27ac ChIP-seq and ATAC-seq for measuring enhancer activity, we also tested taking the geometric mean of DNase-seq and H3K27ac ChIP-seq reads, as proposed by Fulco et al. (2019). For measuring the enhancer-gene contact we evaluated a K562 Hi-C matrix (Rao et al., 2014), an average Hi-C matrix based on 10 different cell types provided by Fulco et al. (2019), and a contact estimate based on a fractal globule model (Lieberman-Aiden et al., 2009). With the fractal globule model the contact can be estimated by the inverse of the distance. We additionally introduced a 5 kb offset for the ABC calculation, resulting in the following contact estimate $c$ at distance $d$:

$$c = max\{d, 5000\}^{-1}\,. \tag{1}$$

We calculated the scores for all genes in the GENCODE hg19 annotation (Frankish et al., 2019), and either used the 5' TSS (ABC, adapted activity, Enformer) or averaged across all TSSs of a gene (avgTSS, gABC). All candidate enhancers within a 10 MB window around the TSS were scored. To construct precision-recall (PR) curves and to calculate the respective area under the curve the score cut-off was incremented by 10,000 steps between the smallest and largest score (Fig. S2; Fig. S3; Tab. S1; Tab. S2; Tab. S4). For each step, the interactions with a score above the cut-off were classified as positive set and compared against the validated

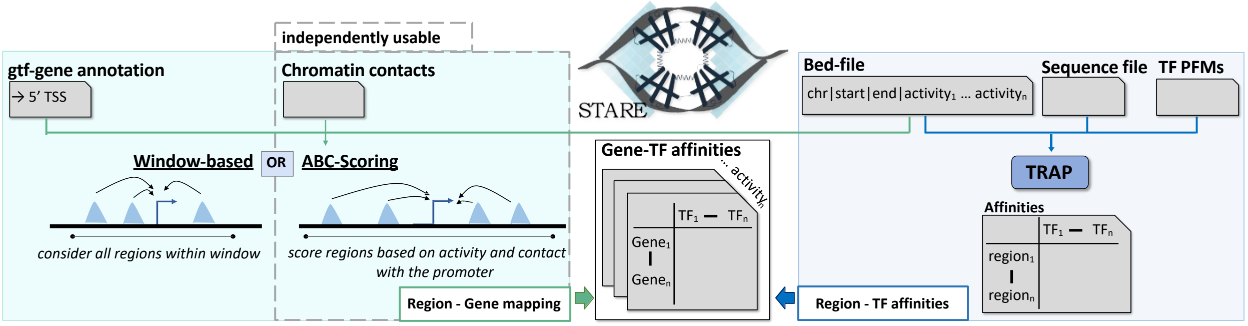


**Figure S1.** Overview of the STARE pipeline. It maps candidate enhancers to their target genes (left panel), either with a window-based approach or an implementation of the ABC-score (Fulco et al., 2019). TRAP (Roider et al., 2007) is used to derive TF binding affinities to candidate enhancer regions (right panel), which are then combined with the enhancer-gene mapping to construct a matrix summarising TF affinities per gene. STARE is adapted to run on multiple cell types with the same candidate enhancers but varying activity, represented by activity columns. PFM: position frequency matrix.

**Table S1.** AUPRC using different ABC-scoring approaches on three CRISPRi-screens with varying assays for measuring enhancer activity and contact frequency. **ABC**: regular ABC-score; **adapted activity**: ABC with gene-specific enhancer activity (main manuscript Eq. 4); **avg TSS**: regular ABC but summarises the activity×contact product across all TSSs of a gene; **gABC**: combination of adapted activity and avgTSS. The highest AUPRC within each validation data set and type of contact data is written in bold.

| Validation data | | Gasperini et al. (2019) 755 valid / 37,738 interactions | | | | Schraivogel et al. (2020) 69 valid / 9,093 interactions | | | | Fulco et al. (2019) 158 valid / 3,999 interactions | | | |
|---|---|---|---|---|---|---|---|---|---|---|---|---|---|
| Enhancer activity assay | | DNase& H3K27ac | DNase | H3K27ac | ATAC | DNase& H3K27ac | DNase | H3K27ac | ATAC | DNase& H3K27ac | DNase | H3K27ac | ATAC |
| K562 Hi-C | ABC | 0.4421 | 0.4333 | 0.3816 | 0.4919 | 0.3600 | 0.3549 | 0.2746 | 0.2909 | 0.4365 | 0.4167 | 0.3835 | 0.3702 |
| | adapted activity | 0.4792 | 0.4795 | 0.4435 | 0.5218 | 0.3775 | 0.3624 | 0.3528 | 0.3228 | 0.4233 | 0.4052 | 0.4030 | 0.3771 |
| | avg TSS | 0.4598 | 0.4562 | 0.3909 | 0.5040 | 0.4861 | 0.4924 | 0.3776 | 0.3976 | 0.4636 | 0.4476 | 0.4030 | 0.3984 |
| | gABC | 0.5042 | 0.5076 | 0.4596 | **0.5408** | **0.5717** | 0.5592 | 0.5050 | 0.4884 | **0.4710** | 0.4552 | 0.4388 | 0.4176 |
| avg Hi-C | ABC | 0.4486 | 0.4395 | 0.3898 | 0.4929 | 0.3554 | 0.3661 | 0.2806 | 0.2945 | 0.4360 | 0.4085 | 0.3860 | 0.3663 |
| | gABC | 0.5038 | 0.5072 | 0.4605 | **0.5364** | **0.5552** | 0.5507 | 0.4985 | 0.4750 | **0.4520** | 0.4387 | 0.4259 | 0.4015 |
| Fractal | ABC | 0.4548 | 0.4478 | 0.4084 | 0.5024 | 0.3781 | 0.3898 | 0.3293 | 0.3268 | **0.4101** | 0.3927 | 0.3762 | 0.3595 |
| | gABC | 0.4616 | 0.4657 | 0.4232 | **0.5034** | 0.5220 | **0.5237** | 0.4566 | 0.4268 | 0.4078 | 0.4004 | 0.3881 | 0.3688 |

interactions. If multiple candidate enhancers overlapped a validated perturbed region, we took the sum of the individual enhancers' score. In addition to PR curves, we calculated the absolute Pearson correlation coefficient between the ABC-/gABC-score and the expression change measured upon perturbation in the CRISPRi-screens (Tab. S3).

To test, whether the difference in the area under the receiver operating characteristic (ROC) curves was significant, we used the R package pROC (Robin et al., 2011), more specifically its implementation of the analysis described by DeLong et al. (1988), which leverages the algorithm of Sun and Xu (2014) for efficiency. We ran pairwise comparisons for the ROC curves between ABC and gABC with different activity assays as input (Tab. S5), as well as between gABC and Enformer (all p-values $\leq 0.05$).

#### 1.1 Detailed comparison of ABC and gABC

To understand for which interactions the gABC-score performs better, we focused on the largest CRISPRi-screen from Gasperini et al. (2019) and selected fixed points in the PR curves where the difference between ABC and gABC was high (40% recall and 60% precision; Fig. S3a-c). At 40% recall the gABC score calls less false positive interactions. We compared the relative enhancer activity and the relative gene-specific enhancer activity. Relative meaning that we divided the activity by the summed activity of all other candidate enhancers of a gene:

$$relativeA_r = \frac{A_r}{\sum_{j \in G_r} A_r}, \quad (2)$$

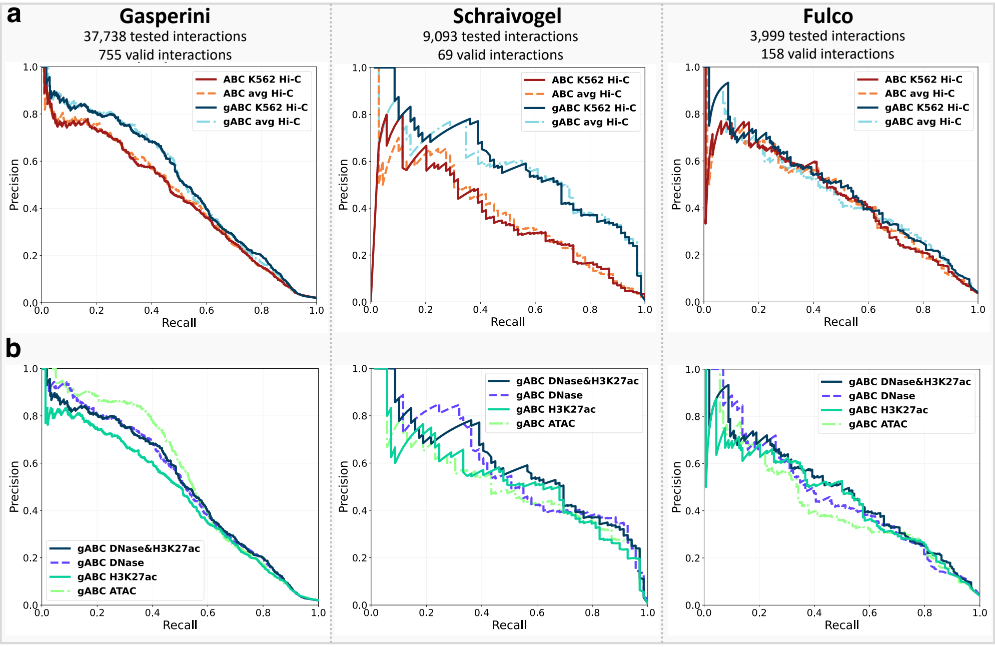


**Figure S2.** Performance of the ABC and gABC-score on three CRISPRi-screens in K562 cells with varying assays for measuring enhancer activity and enhancer-gene contacts. **(a)** Precision-recall (PR) curves of ABC and gABC with K562 Hi-C data (5 kb resolution, Rao et al., 2014) or an average Hi-C matrix. The geometric mean of DNase-seq and H3K27ac ChIP-seq read counts was used as enhancer activity. **(b)** PR curves of gABC with different assays for measuring enhancer activity. The corresponding AUPRC values can be found in Tab. S1. The number of tested and true interactions is the same within columns for (a) and (b).

and in the same manner for the gene-specific activity. We could see that the ABC-score tended to call more false positive interactions for enhancers with higher relative activity (Fig. S3d+e). We also examined the difference on enhancer-level and compared which method calls more false positive target genes for which enhancers at 40% recall. Enhancers with more false positive targets with gABC (n=25) had more potential target genes within their 10 MB scoring window than enhancers for which the ABC-score called more false positives (n=109) (Fig. S3f). We repeated the same comparisons but fixed the score cut-off at 60% precision to describe the true positive interactions called by gABC that ABC missed (n=106), and found that those interactions showed a lower relative enhancer activity (Fig. S3g-i).

#### 1.2 Direct comparison to the implementation of Fulco et al. (2019)

We compared our implementation of the ABC-model directly with the framework of Fulco et al. (2019). The candidate enhancers, their activity, and the chromatin contact data were identical between the two implementations. The main difference was the scoring approach with the gABC-score, as described in the main manuscript. There were also technical differences in deriving the pseudocount for the contact data. We evaluated the performance of both frameworks in the same manner as explained in Sec. 1. The gABC-score reached a higher AUPRC for each combination of input data (Tab. S2). We only included interactions for which we retrieved a score for both implementations, leading to the difference in the number of validated interactions compared to Tab. S1. It should be noted that we did not exclude any interactions where the candidate enhancer overlapped a promoter, nor repressive interactions.

#### 1.3 Comparison to Enformer

For predicting enhancer-gene interactions in K562 cells with Enformer (Avsec et al., 2021) (downloaded from `https://tfhub.dev/deepmind/enformer/1`) we lifted the experimentally validated interactions from Gasperini et al. (2019), Schraivogel et al. (2020) and Fulco et al. (2019) from hg19 to hg38. We kept interactions with a distance of $\leq 96$ kb. This ensures that tested enhancers will always be entirely within the

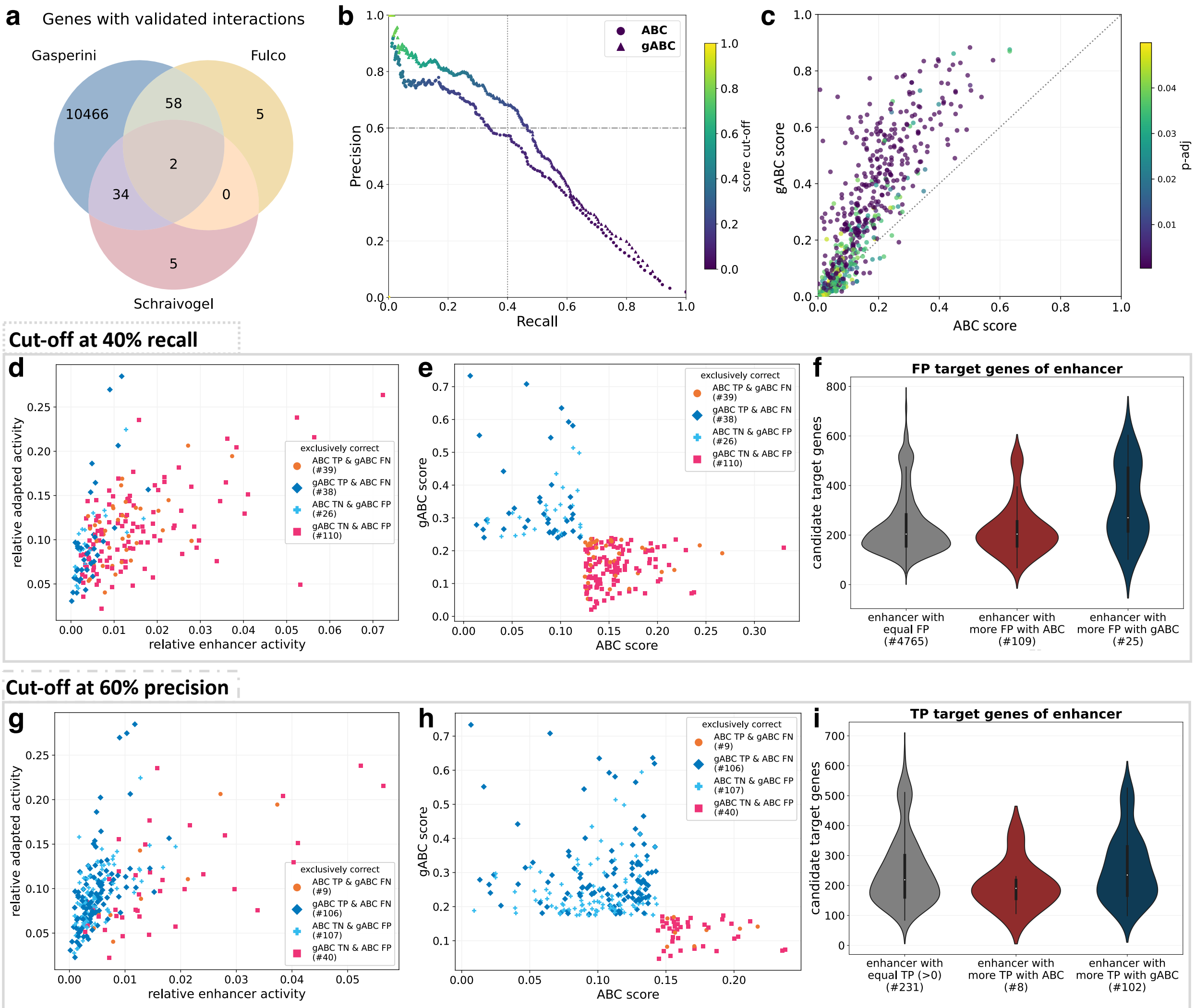


**Figure S3.** Detailed comparison of the ABC- and gABC-score on interactions of the CRISPRi-screen from Gasperini et al. (2019). All shown ABC and g-ABC scores used DNase-seq and H3K27ac ChIP-seq data for measuring enhancer activity and a K562 Hi-C matrix for chromatin contacts. **(a)** Venn diagram of the genes with tested interactions in each CRISPRi-screen. Note that this is a superset of genes with interactions that we could evaluate with the ABC-scores. **(b)** PR curve coloured by ABC/gABC score respectively. The vertical dotted grey line marks the position at 40% recall, the horizontal dash-dotted line 60% precision. **(c)** ABC versus gABC-score of all the significant interactions, coloured by the adjusted p-value. **(d)** Comparison of the relative enhancer activity and relative adapted enhancer activity of interactions where only ABC *or* gABC gives the correct prediction at 40% recall. Relative means the fraction in relation to all other candidate enhancers of the target gene (Eq. 2). *TP*: true positive; *FP*: false positive; *TN*: true negative; *FN*: false negative. **(e)** Showing the same interactions as (d) but now comparing the ABC versus gABC score. **(f)** Distribution of the number of genes within 5 MB range of enhancers, separated by the number of FP target genes called by each method at 40% recall. 'equal FP' contains all enhancer where the number of FP target genes is the same for both scores (0 FP included). 'more FP' means that either score called more FP target genes for that enhancer. The number of enhancers in each category is written on the x-axis. **(g)** Comparison of the relative enhancer activity and relative adapted enhancer activity of interactions where only ABC *or* gABC give the correct prediction at 60% precision. **(h)** Showing the same interactions as (g) but now comparing the ABC versus gABC score. **(i)** Distribution of the number of genes within 5 MB range of enhancers, separated by the number of TP target genes called by each method at 60% precision. 'equal TP' contains all enhancer where the number of TP target genes is the same for both scores (0 TP excluded). 'more TP' means that either score called more TP target genes for that enhancer.

**Table S2.** AUPRC of the original Fulco et al. (2019) pipeline and our gABC-implementation for identifying regulatory interactions on three validation data sets, using varying input assays for measuring enhancer activity. The highest AUPRC within each validation data set is written in bold.

| Validation data | Gasperini et al. (2019) 731 valid / 33,023 interactions | | Schraivogel et al. (2020) 69 valid / 9,058 interactions | | Fulco et al. (2019) 155 valid / 3,996 interactions | |
|---|---|---|---|---|---|---|
| Enhancer activity assay | DNase & H3K27ac | DNase | DNase & H3K27ac | DNase | DNase & H3K27ac | DNase |
| Fulco's pipeline | 0.477 | 0.463 | 0.415 | 0.409 | 0.448 | 0.438 |
| gABC | 0.532 | **0.533** | **0.572** | 0.559 | **0.466** | 0.453 |

**Table S3.** Absolute Spearman correlation coefficient of the ABC- and gABC-score with the absolute measured gene expression change in each CRISPRi-screen. The following columns from CRISPRi-screen data were used for the correlation: Gasperini et al. (2019) "fold_change.transcript_remaining"; Schraivogel et al. (2020) "manual_lfc"; Fulco et al. (2019) "Fraction change in gene expr".

| Validation data | Gasperini et al. (2019) 755 valid / 37,738 interactions | Schraivogel et al. (2020) 69 valid / 9,093 interactions | Fulco et al. (2019) 158 valid / 3,999 interactions |
|---|---|---|---|
| ABC | 0.4663 | 0.4076 | 0.4384 |
| gABC | 0.5425 | 0.4897 | 0.4950 |

**Table S4.** AUPRC of gABC and Enformer on three CRISPRi-screens in K562 cells, limited to interactions with a distance $\leq 96$ kb. For Enformer two different ways were tested to perturb the enhancer sequence in silico, by shuffling the enhancer sequence and by replacing the sequence with neutral nucleotides. The resulting expression differences between native and perturbed sequence were once evaluated via subtraction and once via $log_2$ fold-change.

| Validation data | Gasperini et al. (2019) 633 valid / 5,599 interactions | Schraivogel et al. (2020) 52 valid / 429 interactions | Fulco et al. (2019) 109 valid / 817 interactions |
|---|---|---|---|
| gABC | **0.589** | **0.592** | **0.528** |
| Enformer abs($log_2$FC shuffle) | 0.568 | 0.353 | 0.405 |
| Enformer abs($log_2$FC 'N') | 0.566 | 0.346 | 0.462 |
| Enformer abs(diff shuffle) | 0.534 | 0.273 | 0.359 |
| Enformer abs(diff 'N') | 0.529 | 0.241 | 0.365 |

receptive field of the model. Interactions on the gonosomes and on chr9 were excluded (see Sec. 2). For the interactions from Schraivogel et al. (2020) we only considered interactions to genes which had at least one significant interaction in the whole data set. Enformer was then used to estimate the effect of the CRISPRi perturbation by predicting the CAGE signal at the target gene's 5' TSS using the surrounding 98 kb original sequence as input and comparing it to using an input sequence where the enhancer region was perturbed. The sequence perturbation was done in a 2 kb window centred at the enhancer, once via shuffling (averaged across 25 iterations (Karollus et al., 2022)), and once via replacing the sequence with neutral nucleotides (N). We lifted the interactions back to hg19 and used the difference in the CAGE signal prediction on the native and the perturbed sequence to evaluate Enformer's performance to identify validated interactions from the three CRISPRi-screens. We tested taking the absolute difference, as done by Avsec et al. (2021), and the absolute $log_2$-fold change between CAGE prediction on native and perturbed sequence, for both the shuffling and neutral sequence perturbation approach, under the assumption that removing the sequence of true enhancers should cause a larger change in CAGE signal (Tab. S4). As in Avsec et al. (2021), we averaged Enformer predictions over both strands and small shifts of the input sequence. Avsec et al. (2021) also did a comparison to the ABC-score on the CRISPRi-screens of Gasperini et al. (2019) and Fulco et al. (2019), but calculated the performance binned by distance.

### 2 Processing of chromatin contact data

We used Knight-Ruiz normalisation (Knight and Ruiz, 2012) for all presented contact data. The normalisation did not converge for chr9 of the K562 Hi-C data, likely due to chromosome translocations. Thus, for K562-related data chr9 had to be excluded. During all ABC-score calculations, entries on the diagonal of the contact matrix are replaced by the maximum contact of its neighbouring bins (Fulco et al., 2019), because the contact of a genomic bin with itself is not representative and impaired by measurement artefacts. We add a pseudocount to each enhancer-gene contact, so that all candidate enhancers are taken into account. We model the relation of contact frequency to distance by fitting a linear function to the average contact for each distance $\leq$ 1MB, both in logarithmic space. To compute the average contact we consider all possible contacts, not only the ones $> 0$. We found that excluding the values with 0 contact frequency impairs the linear fit for sparse contact matrices. The pseudocount is adjusted to the distance for interactions $> 1$ MB and set to the estimated contact at 1 MB for interactions $\leq 1$ MB.

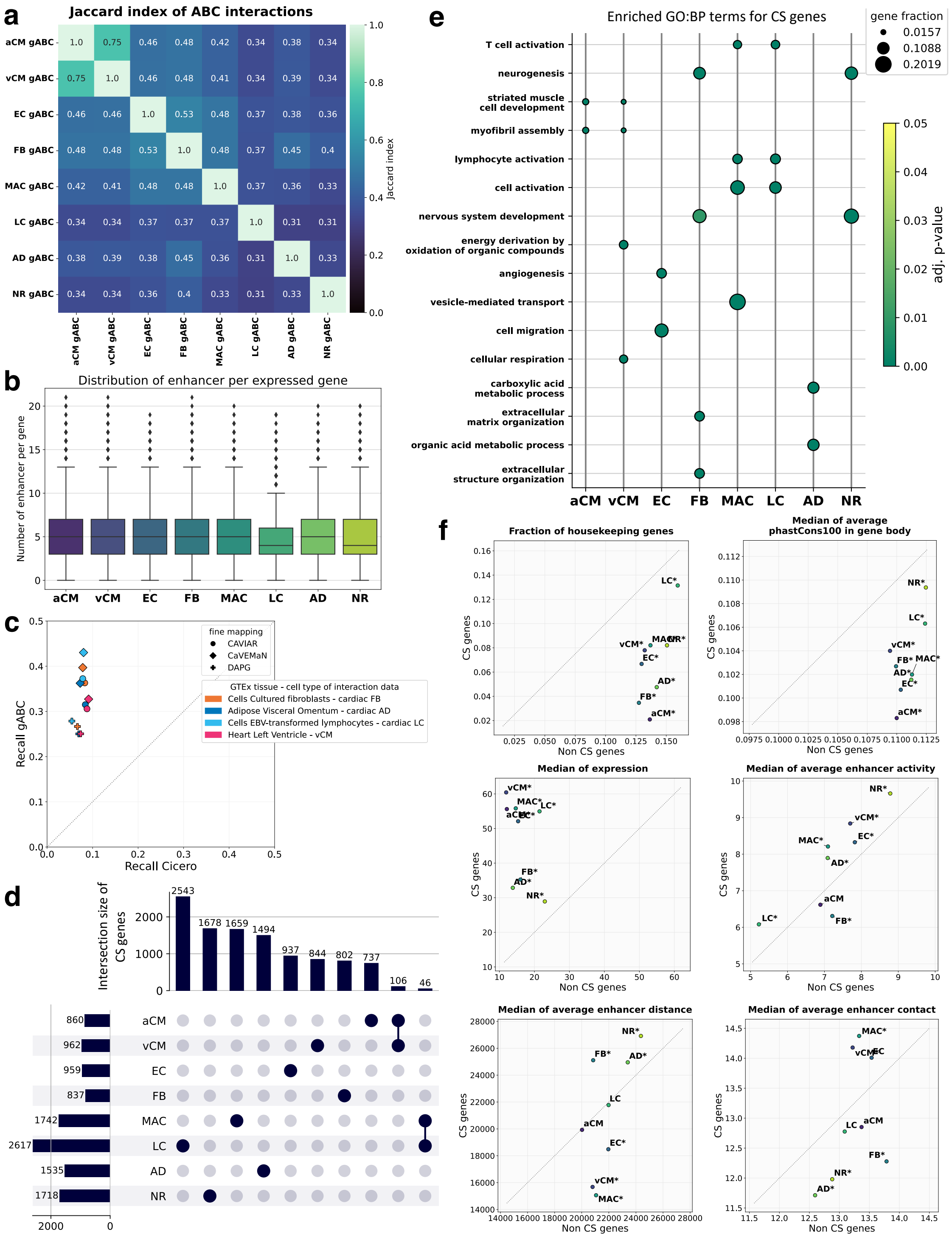


**Figure S4.** Application of STARE on single-cell data (Hocker et al., 2021) and characterisation of cell type-specific (CS) genes (TPM $\geq$ 0.5 and z-score $\geq$ 2). The window size was 5 MB and the gABC score cut-off 0.02. **(a)** Pairwise Jaccard index of the gABC-interactions for each cell type. **(b)** Distribution of the number of enhancers per gene across cell types. **(c)** Intersection of eQTL-gene pairs from different GTEx samples with interactions from a co-accessibility analysis and gABC-interactions. Recall is the fraction of enhancer-gene pairs found by each score out of all pairs where the enhancer contained an eQTL whose target gene was within a distance of 250 kb. The 50,000 highest scored interactions were used, ranked by co-accessibility score or gABC-score respectively. **(d)** Upset plot of CS genes, limited to the 10 largest intersection. **(e)** GO enrichment for CS genes per cell type. Only terms for biological processes (BP) are shown and limited to a maximum of two most significant terms for each cell type. The enrichment analysis was done via g:Profiler (Raudvere et al., 2019). **(f)** Comparison of attributes between CS and not cell type-specific (Non-CS) genes (TPM $\geq$ 0.5). An asterisk indicates a significant difference between CS and Non-CS genes (p-value $\leq$ 0.05). Binary attributes like 'housekeeping genes' were tested with Fisher's exact test, and numeric attributes like the average phastCons100 score with Mann-Whitney U test. The phastCons100 score (Siepel, 2005) was taken as average across the gene body.

**Table S5.** P-values of pairwise tests for the significance of difference of the area under the ROC curves between ABC and gABC using different activity assays as input (DeLong et al., 1988, Robin et al., 2011, Sun and Xu, 2014). The area under the ROC curve was higher for gABC across all comparisons.

| | Gasperini et al. (2019) 755 valid / 37,738 interactions | Schraivogel et al. (2020) 69 valid / 9,093 interactions | Fulco et al. (2019) 158 valid / 3,999 interactions |
|---|---|---|---|
| DNase & H3K27ac | 0.0549 | 0.0014 | 0.0197 |
| DNase | 0.0004 | 0.0044 | 0.0178 |
| H3K27ac | 0.0003 | 0.0001 | 0.0001 |
| ATAC | 0.0405 | 0.0024 | 0.0542 |

### 3 Implementation details of STARE

We combined our implementation of the ABC-scoring with an adaptation of the TEPIC pipeline (Schmidt et al., 2016) and created a new framework, called STARE (Fig. S1). Any ABC-scoring calculation can also be executed independently. STARE was designed under the assumption that cell type-specificity is mainly driven by enhancer activity. Thus, for analysing single-cell data it would be sufficient to define candidate enhancers and measure their activity in individual cells, or summarising activity over clusters of cells or cell types. We call these activity representations activity columns, one for each cell unit. A cell unit can represent any level of aggregation: from individual single cells to metacells up to summarised cell types. STARE was built to leverage this concept of shared candidate enhancers with multiple activity columns. It produces individual results for each activity column, but runs the calculations which are independent of enhancer activity only once. The same holds true for its ABC-scoring implementation. STARE consists of C++ programs connected by a main shell script, where runtime is reduced by utilising parallel computing.

### 4 Application of STARE on single-cell data

We applied STARE on a single-cell data set of the human heart (Hocker et al., 2021) and compared the predicted regulatory enhancer-gene interactions across eight cell types (Fig. S4a). For the expressed genes ($\text{TPM} \geq 0.5$) we found an average of median number of enhancers across cell types of 4.75 ($\text{SD} \approx 0.43$) (Fig. S4b). Based on the snRNA-seq from Hocker et al. (2021) we defined a set of CS genes for each cell type (z-score on expression over all cell types $\geq 2$ and $\text{TPM} \geq 0.5$). Due to the high z-score cut-off, the CS genes were mostly unique to a cell type (Fig. S4d). Lymphocytes (LC) had the largest set with 2,617 CS genes, while fibroblasts (FB) had the smallest set with a size of 837. The two types of cardiomyocytes formed the largest intersection of CS genes shared by more than one cell type. GO term enrichment on the CS genes resulted in terms which appear to fit to the respective cell types, like 'lymphocyte activation' for macrophages (MAC) and LC, or 'extracellular structure organization' for FB (Fig. S4e). To further characterise the sets of CS genes, we examined additional attributes such as conservation or number of enhancers and compared CS genes versus Non-CS genes ($\text{TPM} \geq 0.5$) (Fig. S4f). Across all cell types CS genes were depleted of housekeeping genes. We found that the Non-CS genes were more conserved, as assessed by the average phastCons100 score in the gene body (Siepel, 2005). Additionally, we examined attributes related to the enhancer-gene interactions called by the gABC-score. CS genes had more assigned enhancers than Non-CS genes in all cell types except for LC, and most cell types had a higher fraction of unique interactions. Enhancer activity was elevated for CS genes in six out of the eight cell types. There was no clear trend for enhancer contact or enhancer distance.

Based on the enhancer-gene interactions from each cell type we constructed TF affinity matrices and trained gene expression prediction models for each cell type. We tested different approaches for constructing the TF affinity matrices and compared the prediction performance, both for a training on all genes with measured expression and for a training on CS genes only (Fig. S5a+b). For the models trained on CS genes, we examined TFs with high absolute regression coefficients, as their affinities were informative for expression prediction (Fig. S5c). We found many TFs with known roles, such as NFIX which regulates murine skeletal muscle regeneration through Myostatin expression (Messina et al., 2010; Rossi et al., 2017). NFATC2 and NFATC3 were strongly associated TFs in cardiomyocytes, and NFAT-calcineurin signalling is relevant in regulating hypertrophic growth response in cardiomyocytes (Molkentin et al., 1998; Wilkins and Molkentin,

2002). CTCF with its role in loop formation (Rao et al., 2014) was predicted to have a repressive influence on expression. While a higher number of regions associated to a gene indicated a stronger expression in some cell types, their average size and distance had negative associations.

##### 4.1 Comparison to co-accessible regions

Hocker et al. (2021) provide their results of a co-accessibility analysis on their snATAC-seq with Cicero (Pliner et al., 2018), limited to a distance of 250 kb. We considered a co-accessible region pair as enhancer-gene interaction if either side overlapped a 400 bp window around any annotated TSS of a gene. To build a predictive model of gene expression for each cell type based on the interactions derived from co-accessibility analysis, we retrieved the activities of the interactions in each cell type from the snATAC-seq data. For constructing the TF affinity matrices we also included all regions within 2.5 kb distance to the 5' TSS, as we did for all other approaches.

##### 4.2 Comparison on eQTL data

We intersected interactions from four different heart cell types (Hocker et al., 2021) and K562 cells with eQTL data of matching samples from the GTEx portal (The GTEx Consortium, 2020). We used high confidence eQTL-gene pairs from three different fine-mapping approaches CAVIAR, CaVEMaN and DAP-G (Hormozdiari et al., 2014, Brown et al., 2017, Wen et al., 2016). We mapped those hg38 variants to hg19 with GTEx's lookup table. ABC and gABC are only able to find eQTL-gene pairs where the variant is located in a candidate enhancer and where the distance is not larger than the selected limit for ABC-interactions (2.5 MB for heart cell types; 5 MB for K562 cells). For each tissue and fine-mapping approach we chose a variable number of highest scored ABC-/gABC-interactions and compared the recall, defined as the fraction of enhancer-gene pairs supported by an eQTL that each method finds. We repeated the same procedure for the interactions identified by co-accessibility analysis on the human heart data, which had a smaller distance limit of 250 kb. Since there was a total of 62,384 co-accessible interactions, we compared the top 10,000, 25,000 and 50,000 interactions ranked by co-accessibility score and compared their recall to the respective number of gABC-interactions (Fig. S4c). gABC recovered significantly more eQTL-supported enhancer-gene pairs for all numbers of top-scored interactions (p-value$\leq$0.05 Wilcoxon signed-rank test).

#### 5 Data sources

The presented results are provided via Zenodo (`https://doi.org/10.5281/zenodo.5841991`). The results were produced with STARE's version 1.0.3.1. All data is in hg19. The validated interactions and predictions for K562 cells of Gasperini et al. (2019) were taken from the 'at-scale' data set (GEO: GSE120861). Interactions from Schraivogel et al. (2020) were kindly provided upon request. Validation data from Fulco et al. (2019) was taken from the original publication (41588_2019_538_MOESM3_ESM.xlsx) and their OSF repository (`https://osf.io/uhnb4/`). We used K562 Hi-C data from Rao et al. (2014) (GEO: GSE63525). Sequencing reads in K562 candidate enhancers for DNase-seq and H3K27ac ChIP-seq were already provided in the work of Fulco et al. (2019). Bam files of ATAC-seq data for K562 cells were retrieved from ENCODE: ENCFF128WZG, ENCFF534DCE, ENCFF077FBI (The ENCODE Project Consortium, 2012; Davis et al., 2018). The Enformer model (Avsec et al., 2021) was downloaded from `https://tfhub.dev/deepmind/enformer/1`. Single-cell data of the human heart was taken from Hocker et al. (2021) (GEO: GSE165839). The H3K27ac HiChIP of the left ventricle was kindly provided by Anene-Nzelu et al. (2020). The co-accessible regions called via Cicero (Pliner et al., 2018) were taken from Hocker et al. (2021). eQTL data originates from the GTEx Portal (The GTEx Consortium, 2020) dbGaP accession number phs000424.v8.p2. For the fine-mapped eQTL-gene pairs we took the file 'CAVIAR_Results_v8_GTEx_LD_HighConfidentVariants.gz' for CAVIAR (Hormozdiari et al., 2014), 'GTEx_v8_finemapping_CaVEMaN.txt.gz' for CaVEMaN (Brown et al., 2017) and 'GTEx_v8_finemapping_DAPG.CS95.txt.gz' for DAP-G (Wen et al., 2016). The hg38 variants were mapped to hg19 with the GTEx lookup table 'GTEx_Analysis_2017-06-05_v8_WholeGenomeSeq_838Indiv_Analysis_Freeze.lookup_table.txt.g
The TF motifs can be found in STARE's GitHub repository: `https://github.com/SchulzLab/STARE/blob/`

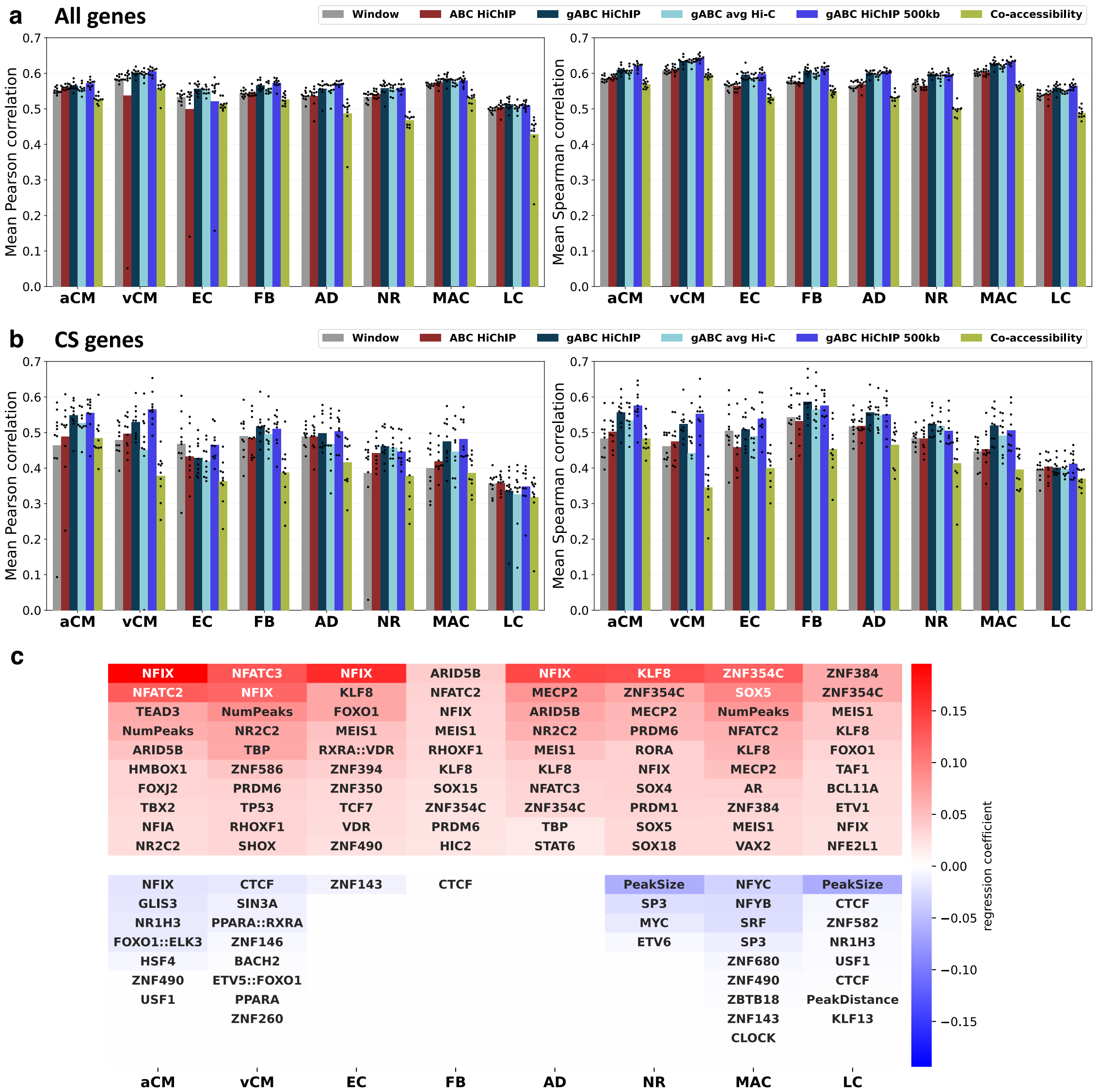


**Figure S5.** Performance of a gene expression prediction model trained on gene-TF affinity matrices based on varying approaches. **(a+b)** The Pearson and Spearman correlation coefficients are shown as average over a 10-fold outer cross-validation. snATAC-seq data was used for all approaches. Window: including all regions within a 5 MB window around the TSS; ABC/gABC H3K27ac HiChIP: regular/generalised ABC-scoring with H3K27ac HiChIP as contact data in a 5MB window; gABC avg Hi-C: gABC with an average Hi-C matrix as contact data in a 5 MB window; gABC H3K27ac HiChIP 500 kb: gABC-scoring with H3K27ac HiChIP as contact data in a 500 kb window; Co-accessibility: enhancer-gene links defined by Cicero Pliner et al. (2018), with a maximum distance of 250 kb. For **(a)** training was done on all genes with an expression value, for **(b)** only on CS genes. **(c)** TFs with the highest absolute regression coefficients in an expression prediction model on CS genes only based on gABC HiChIP. Motif variant suffixes are not shown, different TF motifs of the same TF can have positive and negative correlations in the same cell. **NumPeaks**: number of enhancer considered relevant for a gene; **PeakSize**: average base pair length of the regions assigned to a gene; **PeakDistance**: average distance of regions to the TSS.

`main/PWMs/2.2/Jaspar_Hocomoco_Kellis_human_transfac.txt`. The annotation of housekeeping genes was derived from the HRT Atlas v1.0 (Hounkpe et al., 2021). For the phastCons100 (Siepel, 2005) score we used the track provided by the UCSC genome browser (Navarro Gonzalez et al., 2021).
